# Peripherally-induced regulatory T cells contribute to the control of autoimmune diabetes

**DOI:** 10.1101/199646

**Authors:** Cornelia Schuster, Fangzhu Zhao, Stephan Kissler

## Abstract

Type 1 diabetes (T1D) results from the autoimmune destruction of pancreatic beta cells and is partly caused by deficiencies in the Foxp3^+^ regulatory T cell (Treg) compartment. Conversely, therapies that increase Treg function can prevent autoimmune diabetes in animal models. The majority of Tregs develop in the thymus (tTregs), but a proportion of Foxp3^+^ Tregs is generated in the periphery (pTregs) from Foxp3^-^CD4^+^ T cell precursors. Whether pTregs play a distinct role in T1D has not yet been explored. We report here that pTregs are a key modifier of disease in the nonobesed diabetic (NOD) mouse model for T1D. We generated NOD mice deficient for the *Foxp3* enhancer CNS1 involved in pTreg induction. We show that CNS1 knockout decreased the frequency of pTregs and increased the risk of diabetes. Our results show that pTregs fulfill an important non-redundant function in the prevention of beta cell autoimmunity that causes T1D.

## Introduction

Type 1 diabetes (T1D) is an autoimmune disease in which autoreactive T cells destroy pancreatic beta cells, causing insulin deficiency. The T cells that drive pathogenesis can be held in check by Foxp3^+^ regulatory T cells (Tregs) [1–4]. Several of the gene variants that increase the risk of T1D do so by diminishing the functionality of the Treg compartment. For example, gene variants in the IL-2 pathway that controls the frequency of circulating Tregs have been shown to associate with disease in mouse and human [5–7]. Therapeutic administration of low-dose IL-2 increases the frequency of Tregs and prevents autoimmune diabetes in mouse models[2,4]. This approach is now in clinical trial for the prevention of T1D [8]. The success of low-dose IL-2 therapy in mice illustrates how manipulating the Treg compartment is a promising approach for the treatment of T1D. Yet, we do not fully understand the involvement of different Treg populations in the control of autoimmunity. The majority of Tregs emerge as a distinct population during thymic T cell maturation. Defects in thymic Treg development are a possible cause for the immune dysregulation that leads to autoimmune diabetes [9–11]. However, a proportion of mature Tregs forms not in the thymus but in the periphery [12,13]. These Tregs, termed peripherally-induced or pTregs, derive from mature Foxp3^-^CD4^+^ T cells following the encounter with tolerogenic stimuli, including TGF-β. The highest frequency of pTregs is found in the gut, likely owing to the capacity of gut microbes to promote pTreg induction [14–16]. pTregs are also present in other organs, potentially contributing to systemic immune regulation, but to date, a role for pTregs has only been demonstrated in the gut [17–19], in the lungs [17] and in fetal-maternal tolerance [20]. Whether pTregs participate in immune regulation in other organs is unclear. We set out to ask if pTregs have any role in pancreatic autoimmunity in the context of the nonobese diabetic (NOD) mouse model for T1D. Here, we report that the ability to generate pTregs is a critical modifier of autoimmune diabetes.

## Results

### pTregs are present in the pancreas of NOD mice

Two cellular markers have been proposed to distinguish thymus-derived Foxp3^+^CD4^+^ Tregs (tTregs) from pTregs. Helios and Neuropilin-1 (Nrp1) are both expressed at higher levels in tTregs than pTregs [22–24]. Preliminary measurements with Helios and Nrp1 suggested that the two markers were relatively well correlated, and we opted to focus on Helios because it has been suggested that pTregs may upregulate Nrp1 in inflamed tissues including the pancreas [13]. As reported previously, gut-associated lymphocytes harbored the highest frequency of pTregs. In the colonic lamina propria, 25-30% of all CD4^+^ T cells were Foxp3^+^ pTregs with low levels of Helios (Fig. 1). In contrast, pTregs were rare in the spleen. We detected a slightly higher frequency (~ 0.4% of CD4^+^ cells) in the mesenteric lymph node (mLN) closely associated with the gut. Interestingly, pTregs were also present at similar frequencies in both the pancreatic lymph node (pLN) and in the pancreas (Fig. 1B). The pLN is a major site of activation for autoreactive T cells in autoimmune diabetes. The presence of pTregs in the pLN and in the pancreas suggests that pTregs could affect the autoimmune response that underlies T1D.

**Fig. 1.**
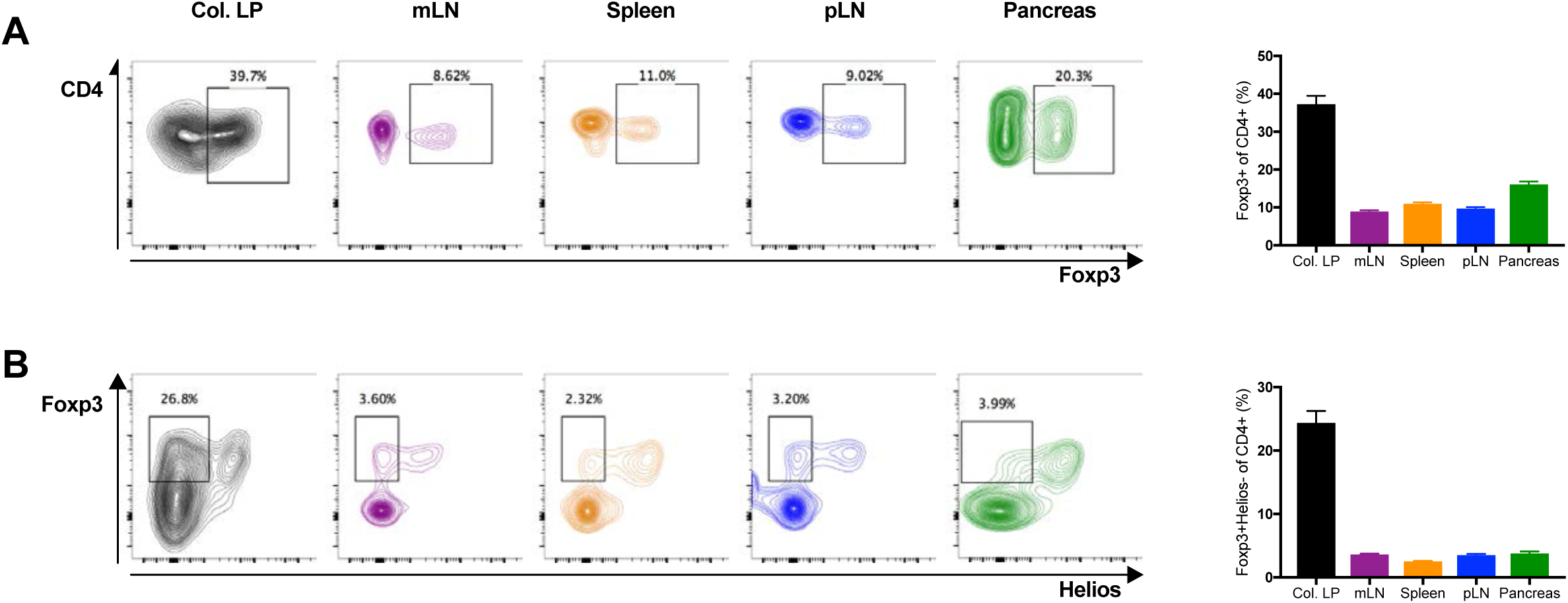
pTregs are present in the pancreas of NOD mice. (A and B) Lymphocytes isolated from the colonic lamina propria (Col. LP in black), mesenteric lymph nodes (mLN in purple), spleen (in orange), pancreatic lymph nodes (pLN in blue) and the pancreas (in green) of NOD WT mice were analyzed by flow cytometry. Total Tregs were characterized as Foxp3^+^ cells (A). pTregs were characterized as Foxp3^+^Helios^-^ cells (B). All samples were gated on live CD4^+^CD8^-^ lymphocytes. Representative FACS plots are shown on the left, cell frequencies (mean ± SEM) on the right. All mice were 6-week old with n=11-25 (3-5 combined individual experiments) per group.

### CNS1 deletion in NOD mice decreases the frequency of pTregs in the pancreas

The differentiation of pTregs involves tolerogenic stimuli that include TGFβ [25,26]. Naïve CD4^+^ T cells exposed to TGFβ during activation adopt a regulatory phenotype. TGFβ stimulation has a direct effect on the induction of Foxp3, the transcription factor characteristic for CD4^+^ Tregs. *De novo* expression of Foxp3 in peripheral CD4^+^ T cells involves the conserved non-coding sequence 1 (CNS1) enhancer in the Foxp3 promoter[21,27], a binding site for Smad3 that acts downstream of TGFβ stimulation [21,28]. CNS1 deletion was shown to decrease the frequency of pTregs [17,27]. CNS1 knockout (KO) is the most specific perturbation available to modify the pTreg compartment without affecting tTregs. We generated CNS1 KO NOD mice using CRISPR-Cas9 genome editing (Fig. 2A). Briefly, we targeted the Cas9 endonuclease to sites immediately upstream and downstream of the CNS1 enhancer that is located at position +2079 relative to the Foxp3 promoter and spans some 630 nucleotides [21]. Two of the mice that developed from microinjected zygotes carried a mutated X-chromosome with the expected CNS1 deletion (Fig. 2B). The CNS1 KO allele was inherited with Mendelian frequency and had no deleterious effects on development or fertility. We measured pTregs in the blood of founder CNS1 KO mice and their progeny, and confirmed that CNS1 deficiency decreased the frequency of pTregs (Fig. 2C and 2D). pTreg frequency was similarly reduced in the colon, spleen and lymph nodes of CNS1 KO NOD mice (Fig. 3B), in accord with published data [27]. Importantly, CNS1 deletion also decreased Helios^-^Foxp3^+^ pTregs in the pLN and in the pancreas (Fig. 3B). Of note, CNS1 KO did not affect the frequency of tTregs or the frequency of total Foxp3^+^ Tregs, except in the colon (Fig. 3A), presumably because the contribution of pTregs to the total Treg population is greatest in this organ. Absolute numbers of CD4^+^ and Foxp3^+^CD4^+^ T cells were comparable in all organs of WT and CNS1 KO mice (not shown). In sum, CNS1 KO decreased pTregs in the gut and in secondary lymphoid organs without affecting tTreg or total Foxp3^+^ Treg numbers overall. Significantly, CNS1 KO diminished the frequency of pTregs in the pancreas and pLN. CNS1 KO NOD mice therefore constitute a new model to specifically investigate the contribution of pTregs to autoimmune diabetes.

**Fig. 2.**
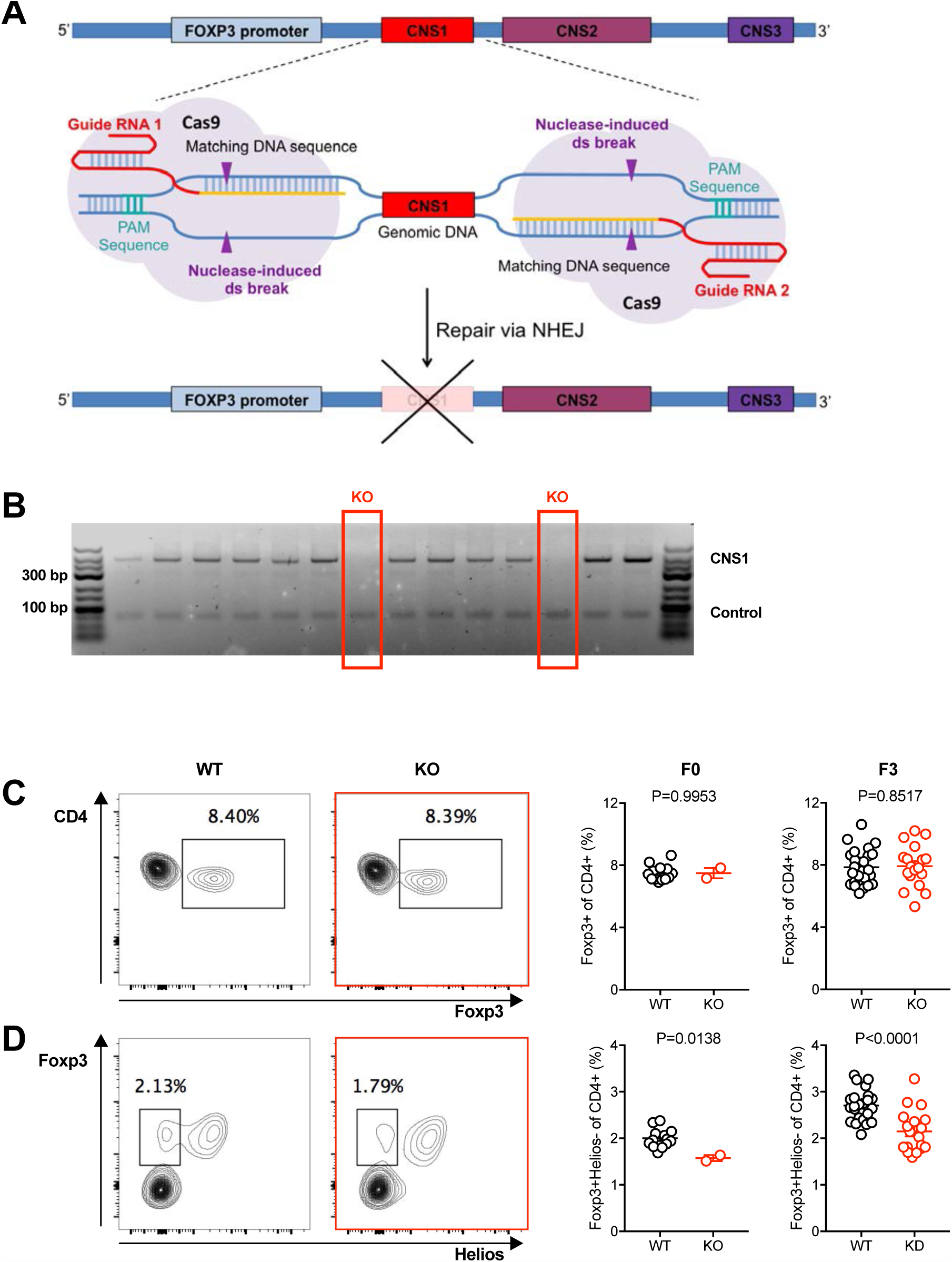
Generation of CNS1 KO NOD mice. (A) Schematic of the strategy used to delete CNS1 using CRISPR/Cas9 genome editing in the Foxp3 promoter region. (B) PCR amplification of the CNS1 region in founder mice to distinguish WT from CNS1 KO mice. (C and D) Blood of founder mice (F0) as well as WT and CNS1 KO littermates in the third off-spring generation (F3) was analyzed by flow cytometry. Total Tregs were characterized as Foxp3^+^ cells (C) and pTregs were characterized as Foxp3^+^Helios^-^ cells (D), all within live CD4^+^CD8^-^ lymphocytes. Representative plots (left panels) and cell frequencies (mean ± SEM, right panels) are shown. All mice were 10-12 weeks old. n=12 (WT), n=2 (CNS1 KO) for F0 and n=25 (WT), n=18 (CNS1 KO) for F3 mice. Exact P-values are shown (two-tailed unpaired t-test).

**Fig. 3.**
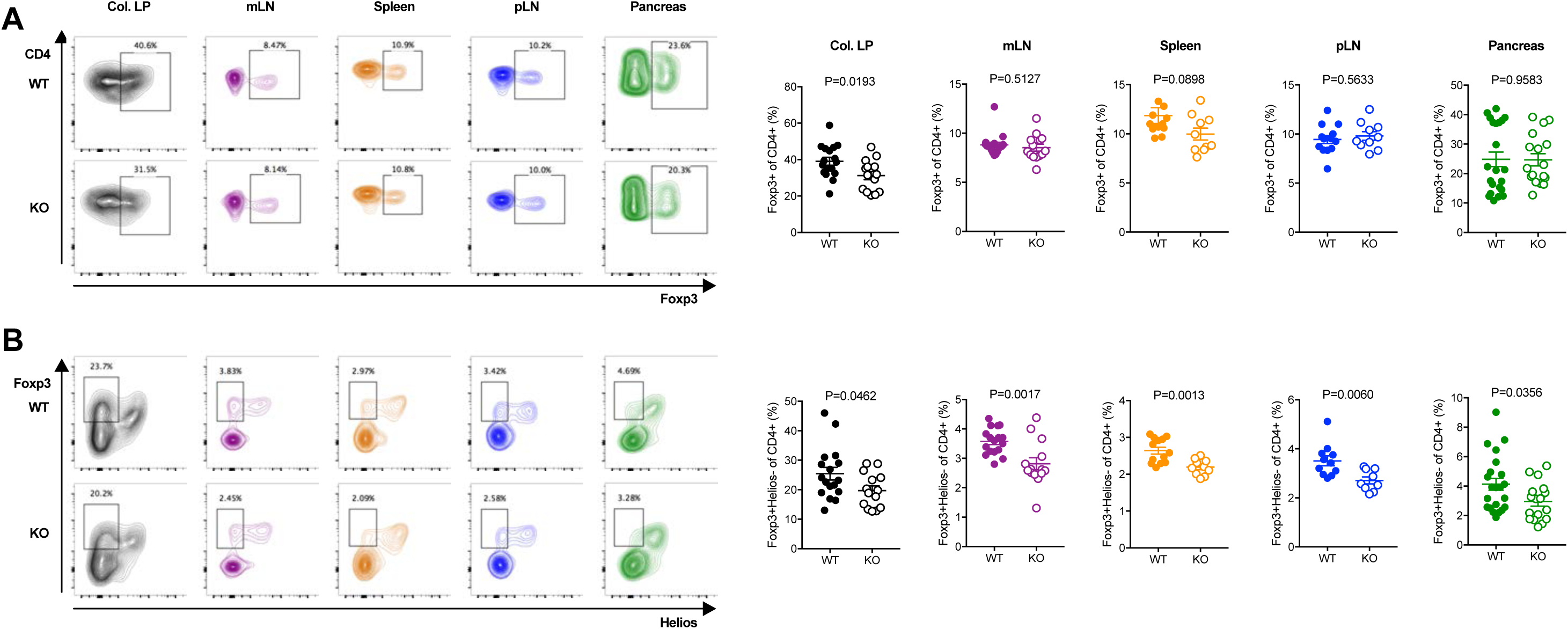
CNS1 KO decreases pTregs in NOD mice. (A and B) Lymphocytes from NOD WT and CNS1 KO colonic lamina propria (Col. LP in black), mesenteric lymph nodes (mLN in purple), spleen (in orange), pancreatic lymph nodes (pLN in blue) and the pancreas (in green) were analyzed by flow cytometry. Total Tregs were characterized as Foxp3^+^ cells (A). pTregs were characterized as Foxp3^+^Helios^-^ cells (B). All samples were gated on live CD4^+^CD8^-^ lymphocytes. Representative FACS plots are shown on the left, cell frequency data (mean ± SEM) on the right. All mice were 6 weeks old, n=9-22 mice per group (3-5 combined individual experiments). Exact P-values are shown (two-tailed unpaired t-test).

### CNS1 deficiency increases the risk of autoimmune diabetes in NOD mice

We bred cohorts of CNS1 KO NOD mice using a breeding scheme that yielded WT and heterozygous (for females only, because Foxp3 is on the X-chromosome) control littermates for comparison. Mice were not separated by genotype at weaning and were co-housed throughout this study. CNS1 KO was reported to cause gastrointestinal pathology in the B6 mouse strain [17], but not in a mixed 129/B6 background [27]. Unlike B6 mice, CNS1 KO NOD mice showed no signs of morbidity or weight loss up to 7 months of age, the period during which mice were monitored for diabetes. We found that CNS1 deficiency increased the frequency of diabetes in both male and female mice (Fig. 4A). Insulitis was more severe in CNS1 KO mice compared with WT littermates (Fig. 4B). We conclude that pTreg deficiency increases the risk of autoimmune diabetes.

**Fig. 4.**
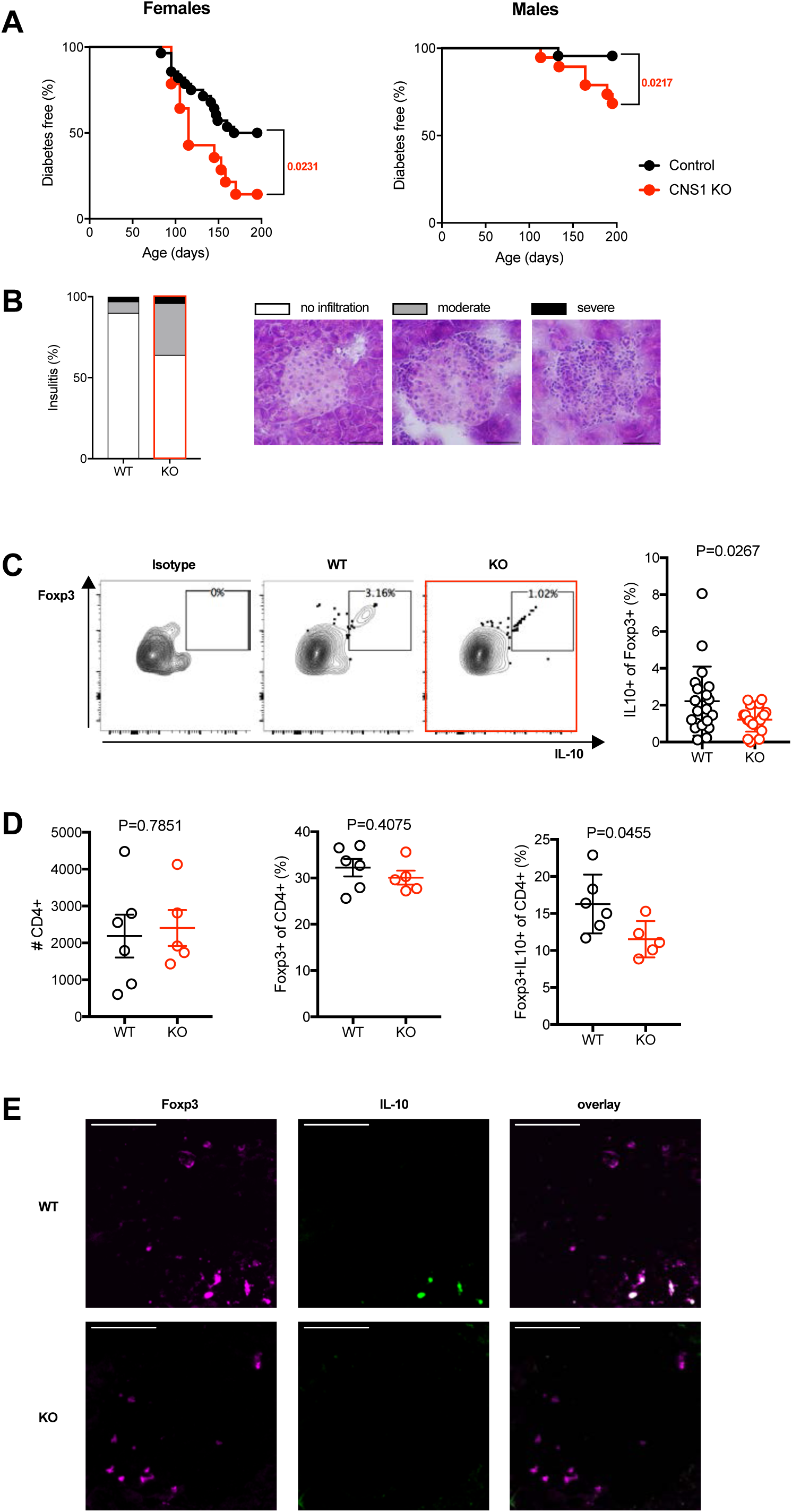
CNS1 KO increases the frequency of diabetes and decreases the frequency of IL-10-producing Tregs in NOD mice. (A) Spontaneous diabetes frequency in WT or heterozygous (Control, n = 28) and CNS1 KO (KO, n = 14) female mice (left) and WT (n=23) and CNS1 KO (n=19) male mice (right). Disease frequency between groups was compared using the Log-rank test, exact P values are shown. (B) Insulitis in female WT (n = 1595 islets) and CNS1 KO (n = 1422 islets) mice at 6 weeks of age. Islets were scored as free of insulitis, or as having moderate or severe infiltration, see representative images (40x magnification, scale bars indicate 50µm). P < 0.0001 (Fisher’s exact test) for the proportion of infiltrated (moderate and severe combined) islets between WT and CNS1 KO groups. (C) Pancreatic lymph node cells from WT (n=20) and CNS1 KO (n=21) NOD mice (all 6 weeks old, 4 combined individual experiments) were stimulated with PMA/ionomycin and analyzed by flow cytometry to measure the frequency of IL-10 producing Foxp3^+^ Tregs. Representative FACS plots including an isotype control antibody for IL-10 are shown on the left, cell frequencies (mean ± SEM) on the right. Values denote frequencies within live CD4^+^CD8^-^Foxp3^+^ lymphocytes. Exact P-values are shown (two-tailed unpaired t-test). (D) Pancreatic lymphocytes from WT (n=6) and CNS1 KO (n=5) NOD mice (all 10 weeks old) were stimulated with PMA/ionomycin and analyzed by flow cytometry to measure the frequency of IL10-producing Foxp3^+^ Tregs. The left panel shows the absolute number of CD4^+^ cells, the middle and right panels show the proportion of total Foxp3^+^ and Foxp3^+^IL10^+^ cells, respectively, within the CD4 population (mean ± SEM). Exact P-values are shown (two-tailed unpaired t-test). (E) Representative pancreas sections from 12-week old WT and CNS1 KO NOD mice stained for Foxp3 (magenta, left panels) and IL-10 (green, middle panels). The overlay (white for co-localization) is shown in the right panels. (40x magnification, scale bar: 50 µm).

### Treg IL-10 production is decreased in the pancreas of CNS1 KO mice

To evaluate the extent of pTreg function in diabetes, we identified IL-10 producing Foxp3^+^CD4^+^ Tregs in the pancreas and pLN. Cells isolated from the pLN (Fig. 4C) and from the pancreas (Fig. 4D) were stimulated overnight then stained for intracellular IL-10. CNS1 KO decreased the frequency of IL-10 producing Foxp3^+^ T cells in both the pLN and pancreas. This was confirmed by immunohistochemical staining that showed a decrease in IL-10^+^Foxp3^+^ cells in the pancreas of CNS1 KO mice (Fig. 4E). Whether decreased IL-10 production itself is causal for increased diabetes frequency in CNS1 KO mice is uncertain, because IL-10 has been ascribed a positive [29–31], negative [32] and neutral [33] role in pancreas autoimmunity by different studies. Notwithstanding, the data show that pTregs impinge on the function of the pancreatic Treg compartment during autoimmune diabetes.

## Discussion

The strongest genetic association with T1D is found in the HLA region [34], implicating T cells as a major driver of autoimmune diabetes. Numerous additional disease-associated genomic regions relate to T cell function [34–36]. In particular, several of the gene variants thought to increase the risk of T1D pertain to the function and homeostasis of Tregs whose defect is thought to be a critical component of T1D pathogenesis [6]. Treg-based therapies are one of the most promising avenues for the treatment of autoimmunity. In this context, it is important that we better understand which Tregs could be leveraged to prevent or halt the autoimmune response that underlies T1D. pTregs may constitute a subset amenable to therapeutic manipulation, owing to the fact that these cells can be induced within the mature T cell population by various agents, including microbial metabolites [37–39]. While inducing pTregs appears feasible, the extent to which these cells contribute to the control of pancreatic autoimmunity had not yet been established. In this report, we showed that pTregs are present in the pancreas and that diminishing their frequency increases the risk of diabetes. Our findings support a role for pTregs in autoimmune diabetes. CNS1 KO was reported to have no effect in the experimental autoimmune encephalomyelitis (EAE) model for multiple sclerosis [17]. A role for pTregs in immune-mediated pathology may thus not be systemic, but has so far only been described in the gut [17–19], the lungs [17] and in fetal-maternal tolerance [20]. The effect of pTregs on islet autoimmunity may relate to the proximity of the pancreas to the gut. The pLN is the primary site of activation for autoreactive T cells [40] in autoimmune diabetes. At the same time, the pLN is readily accessible to antigen and to cells from the gastrointestinal tract [41], providing a direct route by which gut microbe-induced pTregs could impact autoimmunity in T1D. This speculative link would provide a plausible explanation for the effects of the gut microbiota on autoimmune diabetes. It is now well established that diabetes in the NOD mouse model is very sensitive to changes in the microbiome [42–45]. Similarly, prospective studies of human at-risk populations suggest that the gut microbiome modifies the risk of T1D [46,47]. The effects of the microbiome on T1D may derive from the capacity of different microbial communities to promote the generation of pTregs relevant to pancreatic autoimmunity. In light of our finding that pTregs impact the risk of autoimmune diabetes, further exploring this putative mechanism is warranted. Additional research will be required to also determine when and where pTregs affect the autoimmune response that underlies T1D. A pTreg deficit in the gut could modify immune development locally or promote gut leakiness with systemic effects. Alternatively, loss of pTregs in the pLN and in the pancreas could have a direct effect on beta cell autoimmunity. It will also be important to determine at what stage pTregs exert their influence on disease. pTregs may be critical early during immune development when microbial effects on disease risk may be greatest [48]. Alternatively, pTregs may have the capacity to dampen autoimmunity throughout disease development. Notwithstanding the many facets of pTreg function that we do not yet understand, our study demonstrates that pTregs modify the risk of autoimmune diabetes, providing a strong incentive to further explore their role in T1D.

## Materials and Methods

### Mice

CNS1 KO NOD mice were generated by CRISPR/Cas9 genome editing. gRNAs composed of 5’-CACCGAAGACATACACCACCACGG-3’ annealed with 5’-AAACCCGTGGTGGTGTATGTCTTC −3’ and 5’-CACCGCATCAGTCCTCCAGCCAG −3’ annealed with 5’-AAACCTGGCTGGAGGACTGATGC −3’ were cloned into the pX330 vector (Addgene), amplified by PCR and transcribed using the Megashortscript T7 transcription kit (Life Technologies). The two CRISPR target sites are located approximately 120 bp upstream and 40 bp downstream of the CNS1 enhancer that spans 630 bp starting at +2079 bp relative to the Foxp3 promoter [21]. Cas9 mRNA was purchased from Trilink Technologies. RNAs were purified using the Megaclear clean-up kit (Life Technologies). gRNAs and Cas9 mRNA were injected into the pronucleus of NOD zygotes that were reimplanted into pseudo-pregnant Swiss-Webster mice. Genotyping was performed using the PCR primers 5’-GGCGCTTATGTGGCTTCTTTC-3’, 5’-GAGGTAGCTTCTCATTTTCAAGTGG-3’ and 5’-GGAAGCCAACATGGGGTGAA-3’. NOD WT mice were bred and housed in the same room as CNS1 KO mice. All experiments were performed with age- and sex-matched mice and approved by the Institutional Animal Care and Use Committee at the Joslin Diabetes Center.

### Lymphocyte Isolation

Single cell suspensions were prepared from spleen and lymph nodes by mechanical disruption of tissue followed by red blood cell lysis using ACK buffer. Colonic lamina propria cells were isolated after removal of intraepithelial lymphocytes using EDTA followed by digestion with Collagenase Type VIII (Sigma Aldrich) and DNAse (Roche). Cells were collected using a Percoll (GE Healthcare) density gradient. Intra-pancreatic lymphocytes were harvested by digestion using Collagenase P (Roche) followed by a Histopaque-1077 (Sigma) density gradient.

### Intracellular Cytokine Staining

Cells isolated from the pLN and pancreas were stimulated with 50 ng/ml phorbol 12-myristate 13-acetate (PMA, Sigma-Aldrich) and 500 ng/ml ionomycine (Sigma-Aldrich) overnight at 37°C. Golgi-Stop (BD Biosciences) was added for the last 5 h of culture before staining and analysis by flow cytometry.

### Flow Cytometry

Flow cytometry was performed using a LSRII instrument (BD Biosciences). Data were analyzed with the FlowJo software. Fluorescently conjugated CD4, CD8, IL-10, Helios and FoxP3 antibodies were purchased from Biolegend and eBioscience. Dead cells were excluded using the Zombie Aqua^TM^ Fixable Viability Kit (Biolegend). Intracellular staining was performed with a FoxP3-labelling kit (eBioscience).

### Diabetes Frequency Studies

Disease studies were performed with age-matched, contemporary cohorts of mice. Onset of diabetes was monitored by weekly measurements of glycosuria using Diastix (Bayer). Mice with two consecutive readings > 250 mg / dL were considered diabetic.

### Insulitis and Immunohistochemistry

Pancreatic tissue was frozen in OCT (Fisher Scientific) prior to cryosectioning into 7 μm slices. For insulitis scoring, sections were stained with hematoxylin (Fisher Scientific) and counter-stained with eosin (alcoholic Eosin Y, Fisher Scientific). Pancreatic islets were scored as having no infiltration, moderate infiltration or severe infiltration. For immunohistochemistry, pancreas slices were fixed in cold acetone (Sigma-Aldrich), washed twice in PBS and incubated for 1 hour with PBS supplemented with 10% goat serum and 5% Bovine Serum Albumine (Fisher Scientific), then incubated for 1 hour at room temperature with anti-Foxp3 (Cell Signaling Technologies). Slides were washed 3 times in PBS and incubated for 1 hour with a fluorescently labeled secondary antibody (Thermo Fisher Scientific) as well as an Alexa Fluor 488 conjugated IL-10 antibody (Biolegend). All sections were acquired on a Olympus BX-60 microscope equipped with an Olympus DP70 camera using the DPManager software. Fluorescent image analysis was performed with ImageJ software.

### Antibodies

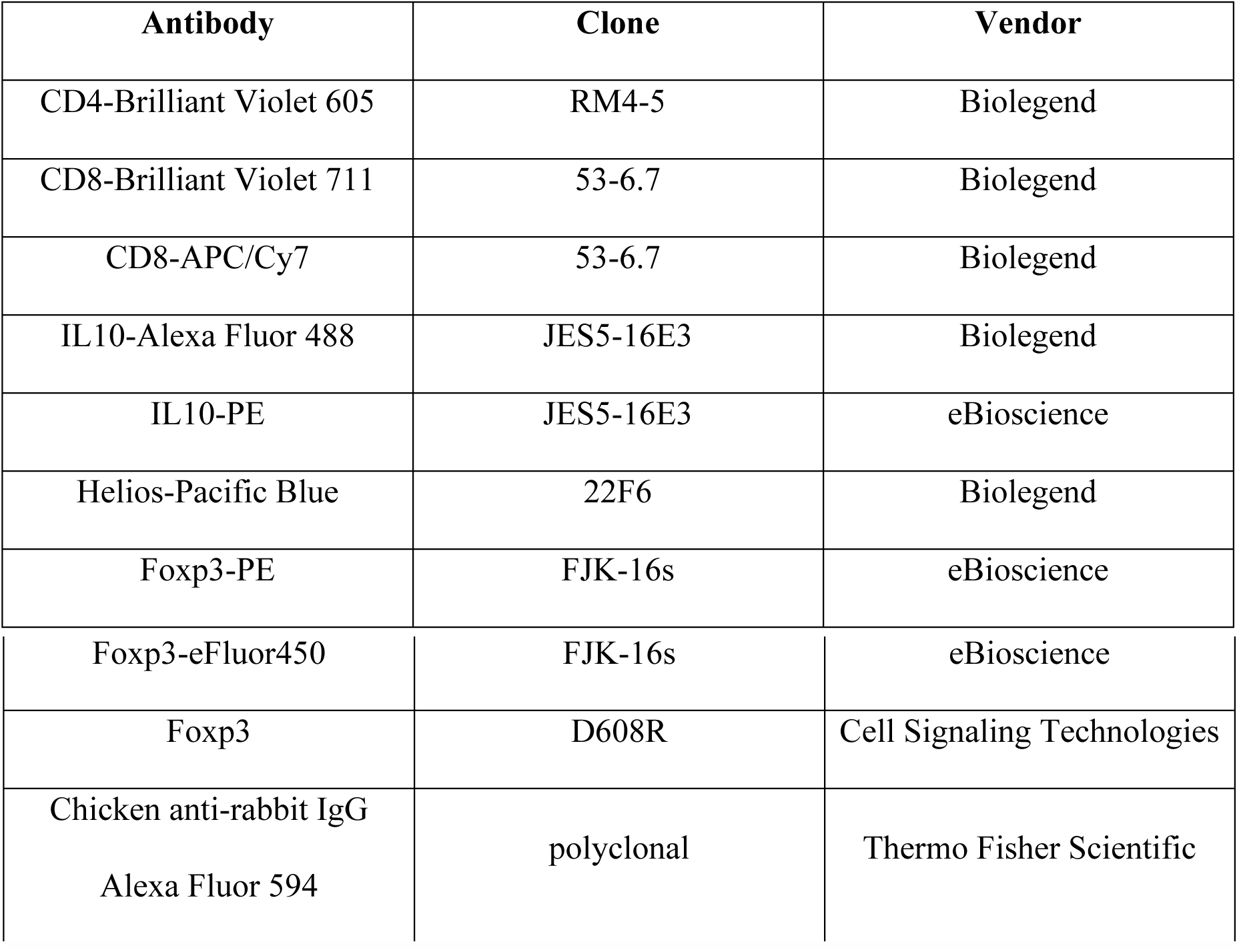

### Statistical Analyses

Data were analysed with the Prism software (Graphpad). Diabetes frequency comparisons were carried out using the Log-rank test. All other comparisons were performed using an unpaired t-test, with P < 0.05 considered significant. Sample sizes were approximated in initial experiments, and adjusted to increase power as needed in replicate experiments.

## Author Contributions

C.S. designed and performed experiments, analyzed data and wrote the manuscript. F.Z. helped with diabetes frequency studies and reviewed the manuscript. S.K. conceived and supervised the project, designed experiments, analyzed data and wrote the manuscript.

## Acknowledgments

The authors wish to thank John Stockton for NOD embryo microinjections and Stephanie Katz for help with mouse colony management. C.S. is the recipient of a postdoctoral fellowship from the Mary K. Iacocca Foundation. This work was supported by NIH funding to the Joslin Diabetes Center (grants P30DK036836 and S10OD021740).

## Conflict of interest disclosure

The authors have no financial conflicts of interest.

